# Sex differences in cytokine induction by activated T cells from hypertensive BPH/2 and normotensive BPN/3 mice

**DOI:** 10.1101/2025.09.30.679479

**Authors:** Devon A. Dattmore, Jeffrey R. Leipprandt, Saamera Awali, Tina Fu, McKenzie Mahlmeister, Hannah Garver, Afolashade Onunkun, D. Adam Lauver, Cheryl E. Rockwell

## Abstract

Over the past two decades, considerable evidence has emerged to implicate a role for the immune system in the development of hypertension. Previous studies have shown immune cells contribute to the development of hypertension in multiple animal models, however the role of the immune system in spontaneously hypertensive BPH/2 mice is not clear. In the current studies, found T cells derived from male hypertensive BPH/2 mice demonstrated an attenuated activation as compared to those derived from male BPN/3 normotensive mice. However, we also observed striking sex differences in T cell cytokine production in these strains. At 24 h post activation, in comparison to male BPH/2 mice, activated T cells from male BPN/3 mice secreted more IL-2, IL-3, IL-4, IL-6, IL-10, IL-17A, IL-17F, IL-22 and TNFα. In contrast to male mice, less than half of these cytokines were different between strains in female mice. We also noted marked differences in early Th17 cytokine production in which IL-17A, IL-17F and IL-22 were greater in the male, but not female, BPN/3 groups. Taken together, the data suggest that polyclonally activated T cells from male, and to a much lesser extent, female BPH/2 mice have a weaker cytokine response as compared to T cells from BPN/3 mice which may be due to an overall attenuated activation of T cells from male BPH/2 mice. Overall, while there are striking differences in T cell response between the BPH2 and BPN/3 strains in male mice, the data indicate far fewer differences between the strains in female mice.

## Introduction

While there are numerous genetic rat models of hypertension, BPH/2 mice are one of the few spontaneously hypertensive mouse models. Generated in the 1960’s and 1970’s by the geneticist, Dr. Gunther Schlager, the hypertensive BPH/2 strain and the normotensive BPN/3 control strain were developed via an 8-way cross of the inbred mouse strains: C57BL/6J, BALB/cJ, LP/J, SJL/J, 129/J, CBA/J, RF/J and BDP/J [1; 2; 3; 4]. Following the 8-way cross, the mice were then stratified by blood pressure and bred within the elevated, low and normal blood pressure groups. The BPH/2 mice are the result of selective breeding of mice with elevated blood pressure and are characterized by a substantial and early rise in systolic and diastolic blood pressure (occurring as early as 6 weeks) in both male and female mice.

Sex differences remain an important factor in hypertension that is pertinent to both human disease and preclinical animal models [5]. With respect to humans, it is well established that there is increased prevalence of hypertension in men under the age of 50 as compared to women. In contrast, women over the age of 50 have greater risk of hypertension compared to men [6]. The correlation of these effects with the onset of menopause suggests hormones may play at least a partial role in the mechanism [7].

Likewise, there are a number of important sex differences in the development of hypertension in animal models [5]. Numerous animal models of experimental hypertension result in higher blood pressure in males compared to females. These include angiotensin II (Ang II)-infused Sprague Dawley rats, Ang II-infused C57Bl/6 mice, spontaneously hypertensive rats (SHR), stroke-prone spontaneously hypertensive rat (SHRSP) and Dahl salt-sensitive rats [8; 9; 10; 11; 12; 13; 14; 15; 16; 17]. The mechanisms behind the sex differences in these various models are complex but likely involve both hormonal as well as Y-linked genetic factors in many cases. The increased incidence and magnitude of hypertension in animal models is often matched by sex differences in inflammation and immunity. For example, the greater increase in blood pressure in Ang II-infused male C57Bl/6 mice correlates with elevated proinflammatory cytokines and decreased NO production [14; 15]. Likewise, Ang II-infused male Sprague Dawley rats show a greater increase in blood pressure as well as increased proinflammatory T cells and decreased numbers of anti-inflammatory regulatory T cells [18].

In comparison to other experimental models of hypertension, previous studies have not shown differences in systolic blood pressure between sexes in the BPH/2 spontaneously hypertensive mouse model [19]. However, whether there are differences in immune cell function between sexes in the hypertensive BPH/2 and normotensive BPN/3 mice is not known and is investigated in the current study.

## Methods

### Materials

The tissue culture-grade reagents: RPMI 1640, Pen/Strep, HEPES, sodium pyruvate and nonessential amino acids were obtained from ThermoFisher Scientific (Waltham, MA). Fetal bovine serum was purchased from Biowest LLC (Kansas City, MO). The anti-CD3 and anti-CD28 antibodies used for splenocyte activation were purchased from BD Biosciences (Franklin, NJ). The Fab2 cross-linking antibody was obtained from Jackson Immunoresearch (West Grove, PA). All other materials were purchased from Millipore Sigma (St. Louis, MO) unless otherwise indicated.

### Animals

All animal protocols and procedures are aligned with the Guide for the Care and Use of Animals and were approved by the Institutional Animal Care and Use Committee at Michigan State University. Five-week-old male and female BPH/2 and BPN/3 mice were purchased from Jackson Laboratories (Bar Harbor, ME). The mice were maintained in specific pathogen-free conditions and were allowed to acclimate for one week prior to initiating studies. Food and water were provided *ad libitum* and the animals were maintained on a 12 h light-dark cycle.

### Blood Pressure Measurements

Mean arterial blood pressure was measured in conscious BPN/3 and BPH/2 mice using tail-cuff plethysmography (CODA® High Throughput system, Kent Scientific). Mice were acclimated to restraint and tail-cuff inflation over several days prior to data collection to minimize stress-related artifacts. Multiple measurements were recorded at each session and averaged for analysis. Surgical implantation of radiotelemetry devices was attempted for continuous blood pressure monitoring. However, implantation was not successful due to the very small size of the female BPH/2 mice.

### Tissue Collection

Spleen and mesenteric perivascular adipose tissue (mPVAT) were collected from 24-week old male and female BPH/2 and BPN/3 mice and processed into single cell suspensions. The splenocytes were immediately pipetted into culture plates and placed into an incubator at 37°C 5% CO_2_ for subsequent activation and analysis.

### Activation of splenic immune cells

Isolated splenocytes were cultured in complete media (RPMI 1640, 10% fetal bovine serum, 100 U/ml penicillin, 100 U/ml streptomycin, 25 mM HEPES, 1 mM sodium pyruvate and 10 mM nonessential amino acids). The cells were cultured in 96-well culture plates and activated with anti-CD3 (1.5 µg/ml), anti-CD28 (1.5 µg/ml) and Fab2 cross-linking antibody (1.5 µg/ml). The cell supernatants were collected at either 24 h or 120 h post activation for subsequent analysis.

### Cytokine Quantification

Cytokine concentrations were quantified in cell supernatants by EVE Technologies (Calgary, Alberta, CA). The samples were analyzed by multiplex bead array using the MD32 and MD12-TH17 panels.

### Statistical Analysis

The mean +/- SEM were calculated for each group. For normally distributed data, the groups were analyzed by 2-way ANOVA followed by Dunnett’s post hoc analysis (GraphPad Prism 10). For non-normally distributed data, the data were first log-transformed prior to analysis by 2-way ANOVA and Dunnett’s post hoc test (GraphPad Prism 10).

## Results

### Elevated blood pressures in male and female BPH/2 mice

Previously published studies have consistently reported elevated blood pressure in BPH/2 mice as compared to BPN/3 mice—an effect observed in both males and females. Consistent with this, we also found higher mean arterial pressure in both male and female BPH/2 mice (Fig. 1). We also found that BPH/2 mice were smaller than BPN/3 mice and had proportionately smaller spleens resulting in no difference in the spleen-to-body weight ratio (Fig. 2). In contrast to spleen, female BPH/2 mice had a greater ratio of mesenteric perivascular adipose tissue (mPVAT) to body weight (Fig. 2). No difference in the ratio of mPVAT-to-body weight was observed between male mice in the two strains.

**Figure 1.**
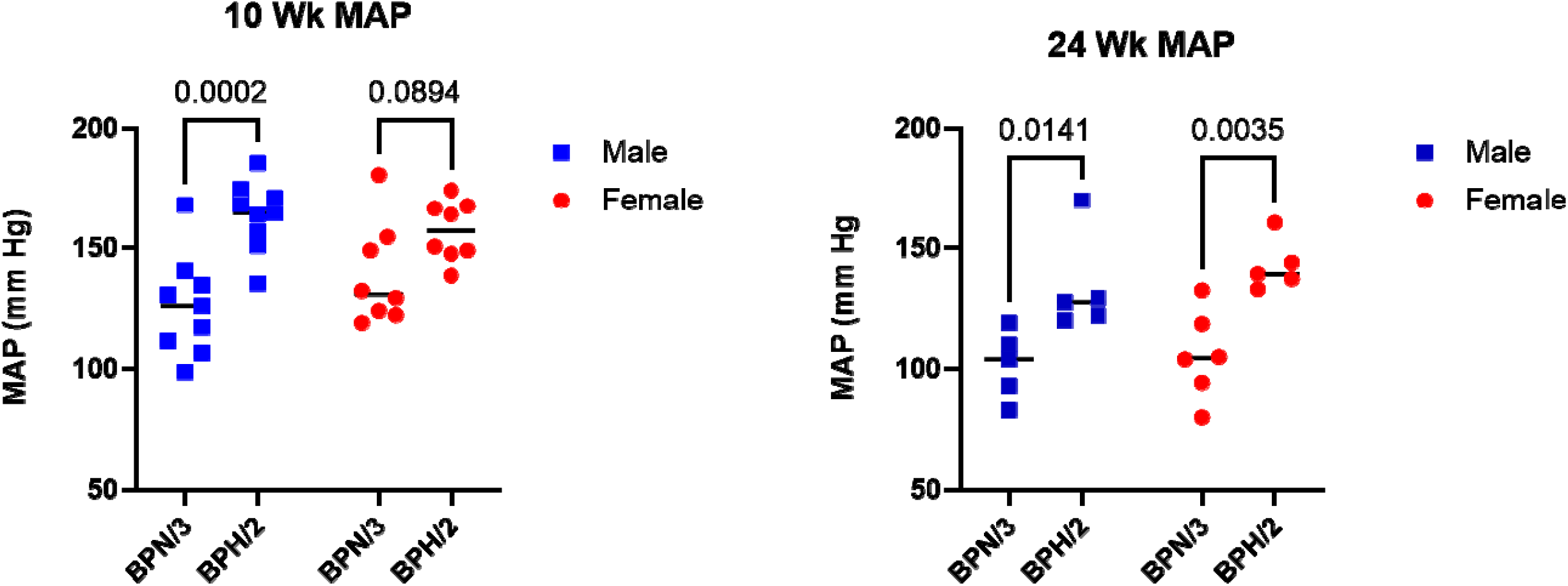
Both male and female BPH/2 mice have elevated arterial blood pressure as compared to BPN/3 mice. Blood pressure was determined by tail cuff plethysmography. Shown in the graphs are the mean arterial pressures for BPN/3 and BPH/2 mice at 10 and 24 wk. Data are presented as scatter plots and include the mean (n = 9 for 10 wk, n = 5 - 6 for 24 wk). p values as determined by two-way ANOVA followed by Dunnett’s post hoc test.

**Figure 2.**
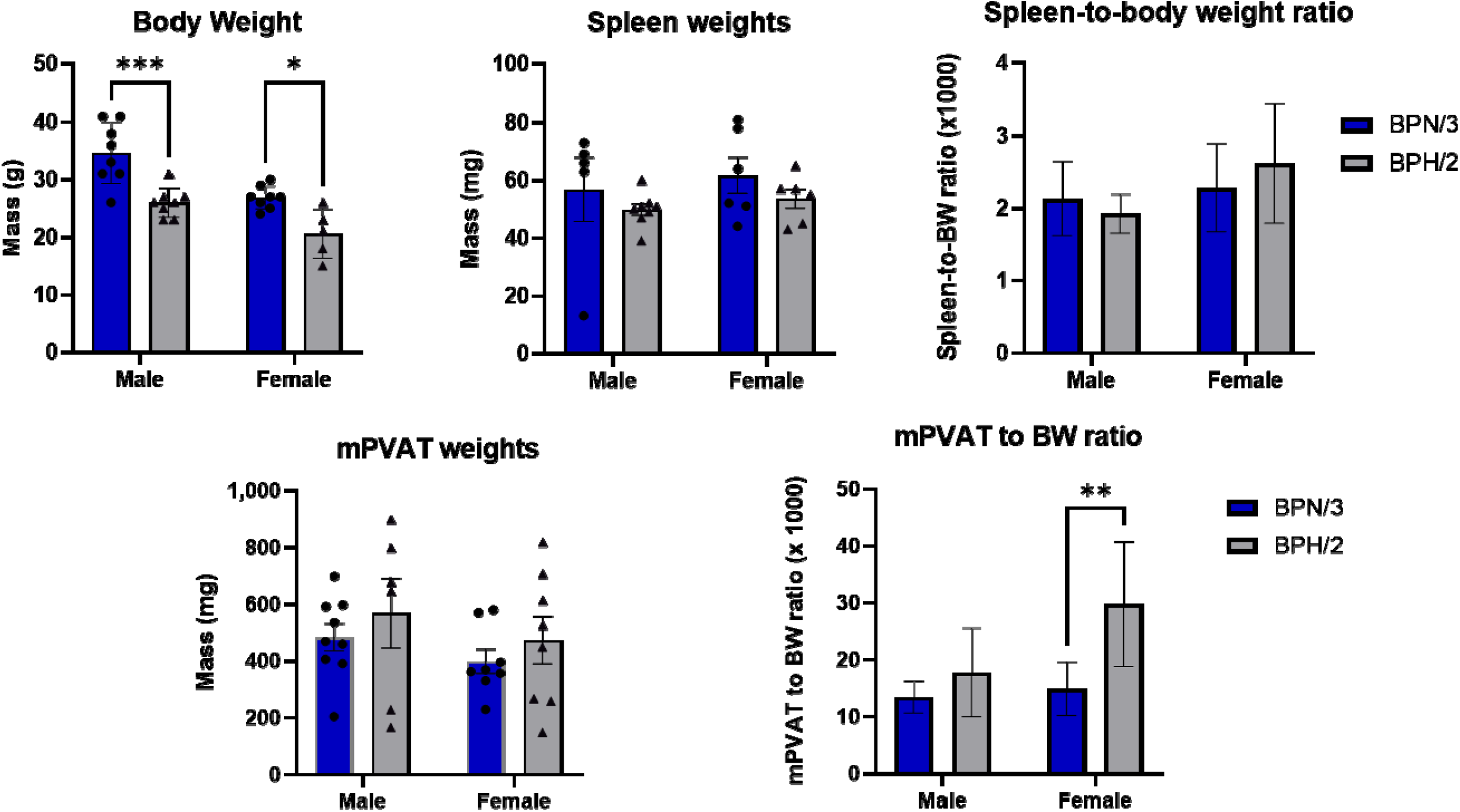
Both male and female BPH/2 mice have lower body weights as compared to BPN/3 mice. Shown in the graphs are the body weights, spleen weights, and spleen-to-body weight ratios, mesenteric perivascular adipose tissue (mPVAT) weights and mPVAT-to-body weight ratios for BPN/3 and BPH/2 mice. Data are presented as bar graphs or scatter plots plus bar graphs and include the mean ± SEM (n = 5). * p<0.05, ** p<0.01 and ***p<0.001 as determined by two-way ANOVA followed by Dunnett’s post hoc test.

### Decreased production of cytokines and chemokines by activated immune cells from male BPH/2 mice

To determine whether there are differences in immune cell function, we quantified cytokine and chemokine secretion by immune cells that were polyclonally activated ex vivo. We found notable differences between both the two strains as well as between the sexes. Specifically, in immune cells activated for 24 h from male BPH/2 mice, we observed significantly decreased secretion of nearly all T cell cytokines (Fig. 3). In contrast to the male mice, there were fewer strain differences in the cells from the female mice. Likewise, we observed a similar trend in chemokine secretion in which cells from male BPH/2 mice secreted significantly lower levels of most chemokines as compared to cells from male BPN/3 mice (Fig. 4). Production of cytokines by activated T cells stimulated secondary secretion of cytokines by myeloid and stromal cells. We found that most of these secondary myeloid/stromal cytokines were significantly lower in the male BPH/2 group as compared to the male BPN/3 group (Fig. 5). In addition to measuring early cytokine expression, we also quantified late cytokine production at 120 h post activation. Here, the effects were similar to those on early cytokines in which cells from male BPH/2 mice produced significantly lower levels of cytokines as compared to those from male BPN/3 mice (Fig. 6). Again, these strain differences were much less pronounced in female mice. Overall, the data show markedly decreased cytokine and chemokine production by immune cells from male, and to a lesser extent female, BPH/2 mice as compared to immune cells from BPN/3 mice.

**Figure 3.**
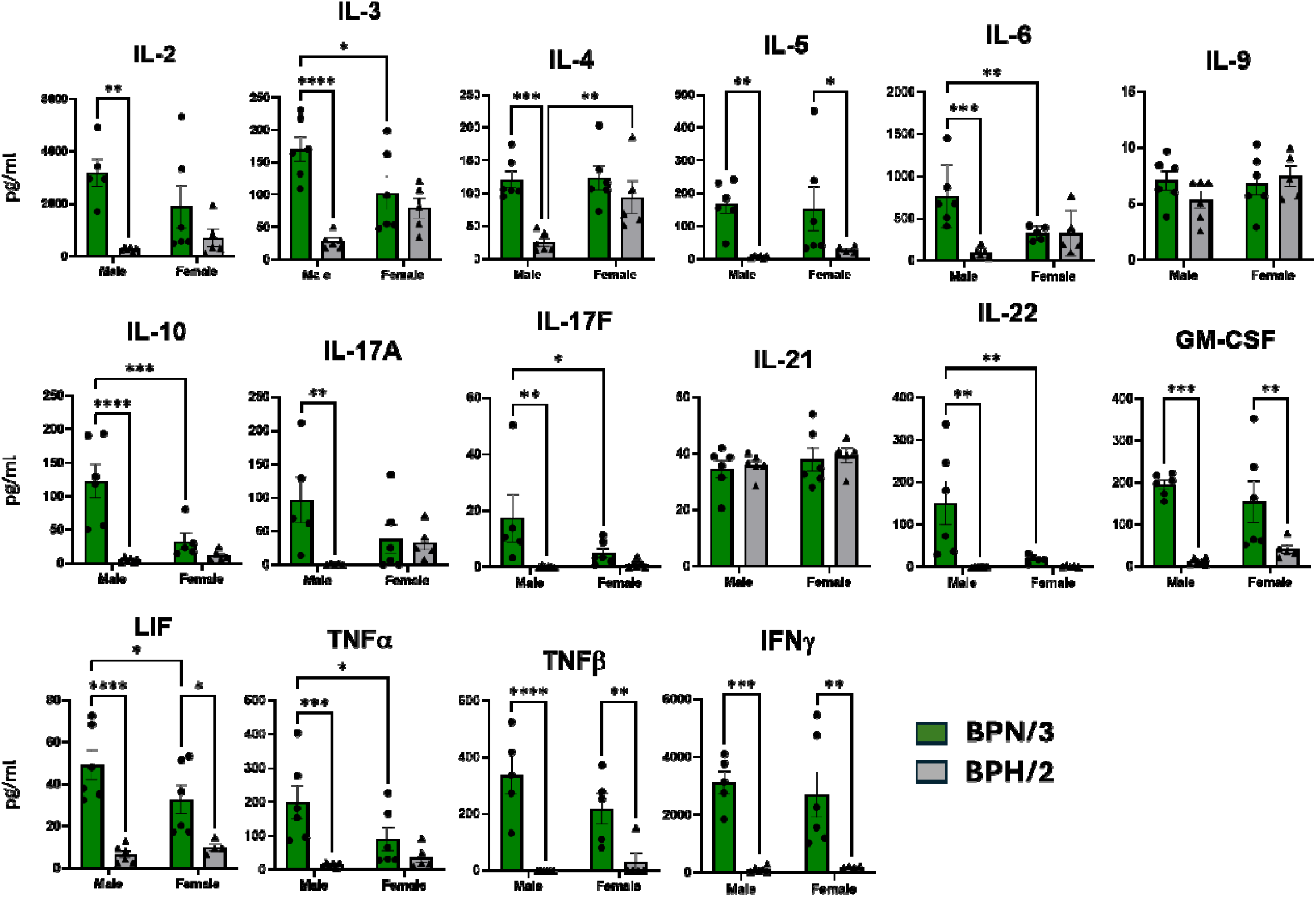
Reduced induction of T cell cytokines by splenic T cells from male BPH/2 mice 24 h after activation. Freshly isolated splenocytes were cultured in the presence or absence anti-CD3/anti-CD28 activating antibodies. Cell supernatants were collected after 24 h for cytokine analysis by EVE Technologies (Calgary, Alberta, Canada). Data are presented as scatter plots plus bar graphs and include the mean ± SEM, n = 6. * p<0.05, ** p<0.01, ***p<0.001 and ****p<0.0001 as determined by 2-way ANOVA followed by Dunnett’s post hoc analysis.

**Figure 4.**
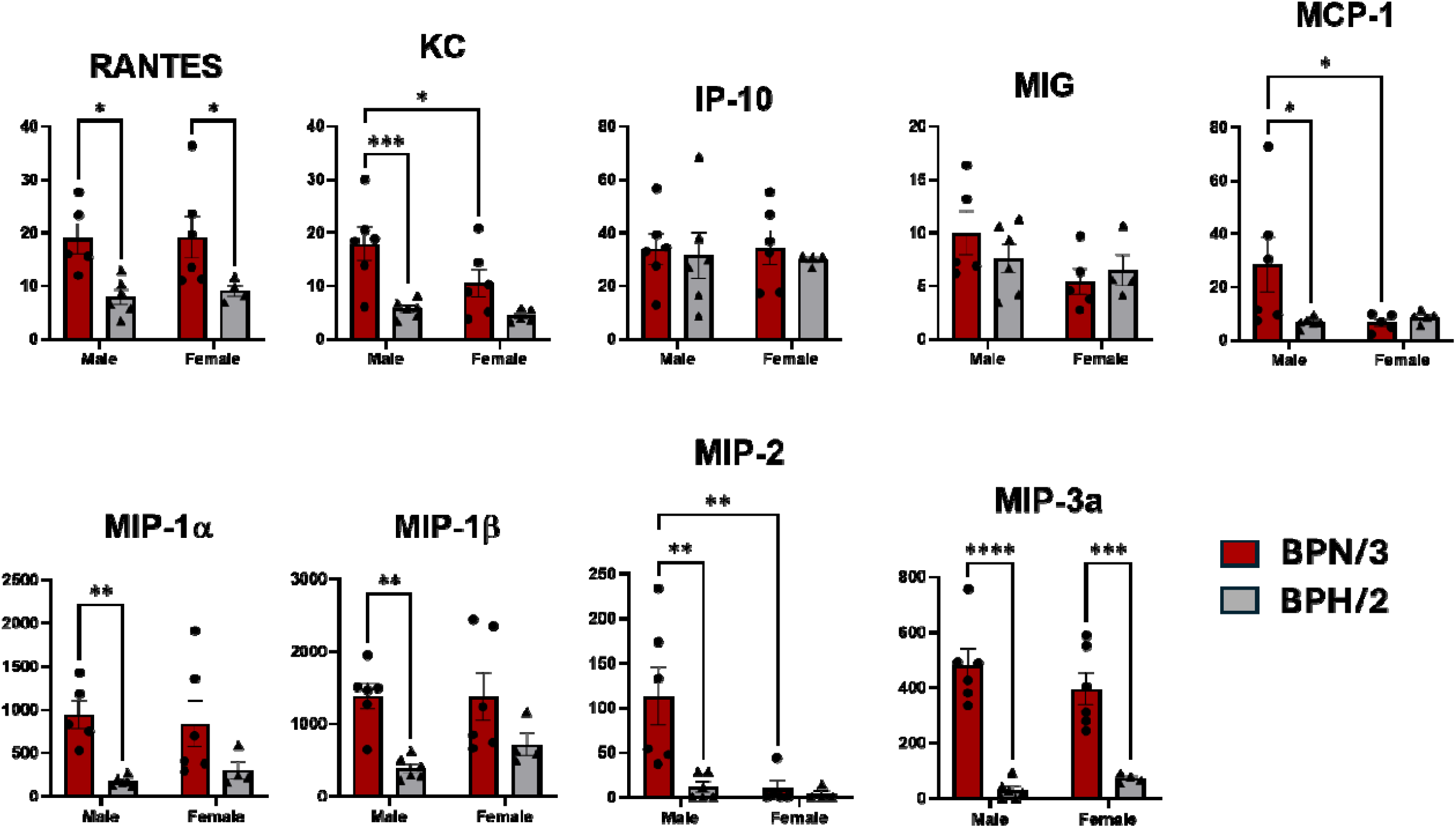
Reduced induction of most chemokines by splenic T cells from male BPH/2 mice 24 h after activation. Freshly isolated splenocytes were cultured in the presence or absence anti-CD3/anti-CD28 activating antibodies. Cell supernatants were collected after 24 h for cytokine analysis by EVE Technologies (Calgary, Alberta, Canada). Data are presented as scatter plots plus bar graphs and include the mean ± SEM, n = 6. * p<0.05, ** p<0.01, ***p<0.001 and ****p<0.0001 as determined by 2-way ANOVA followed by Dunnett’s post hoc analysis.

**Figure 5.**
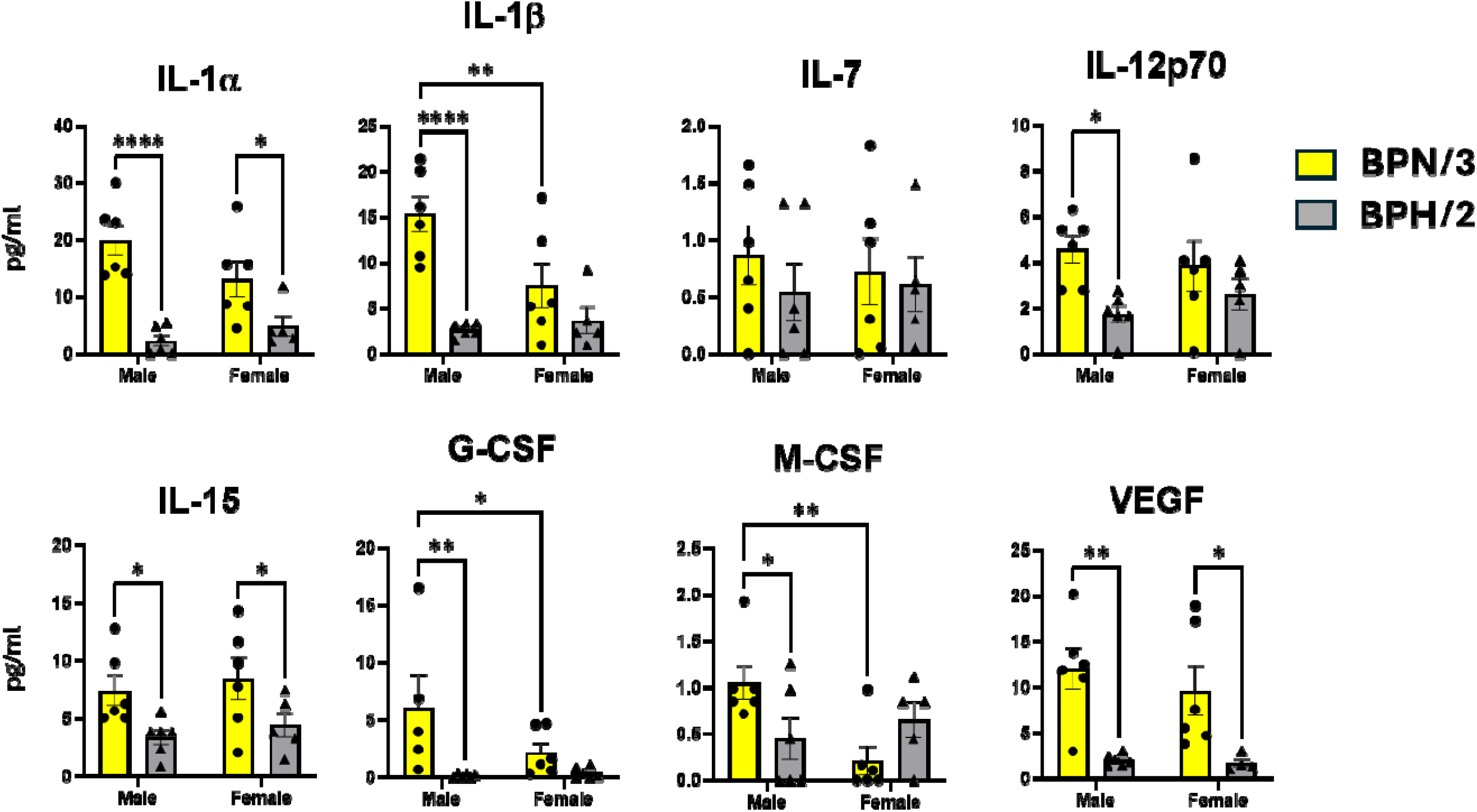
Decreased induction of secondary cytokines by myeloid/stromal cells from male Bph/2 mice 24 h after activation. Freshly isolated splenocytes were cultured in the presence or absence anti-CD3/anti-CD28 activating antibodies. Cell supernatants were collected after 24 h for cytokine analysis by EVE Technologies (Calgary, Alberta, Canada). Induction of cytokines was likely the result of indirect stimulation of myeloid cells by cytokines released by T cells. Data are presented as scatter plots plus bar graphs and include the mean ± SEM, n = 6. * p<0.05, ** p<0.01, ***p<0.001 and ****p<0.0001 as determined by 2-way ANOVA followed by Dunnett’s post hoc analysis.

**Figure 6.**
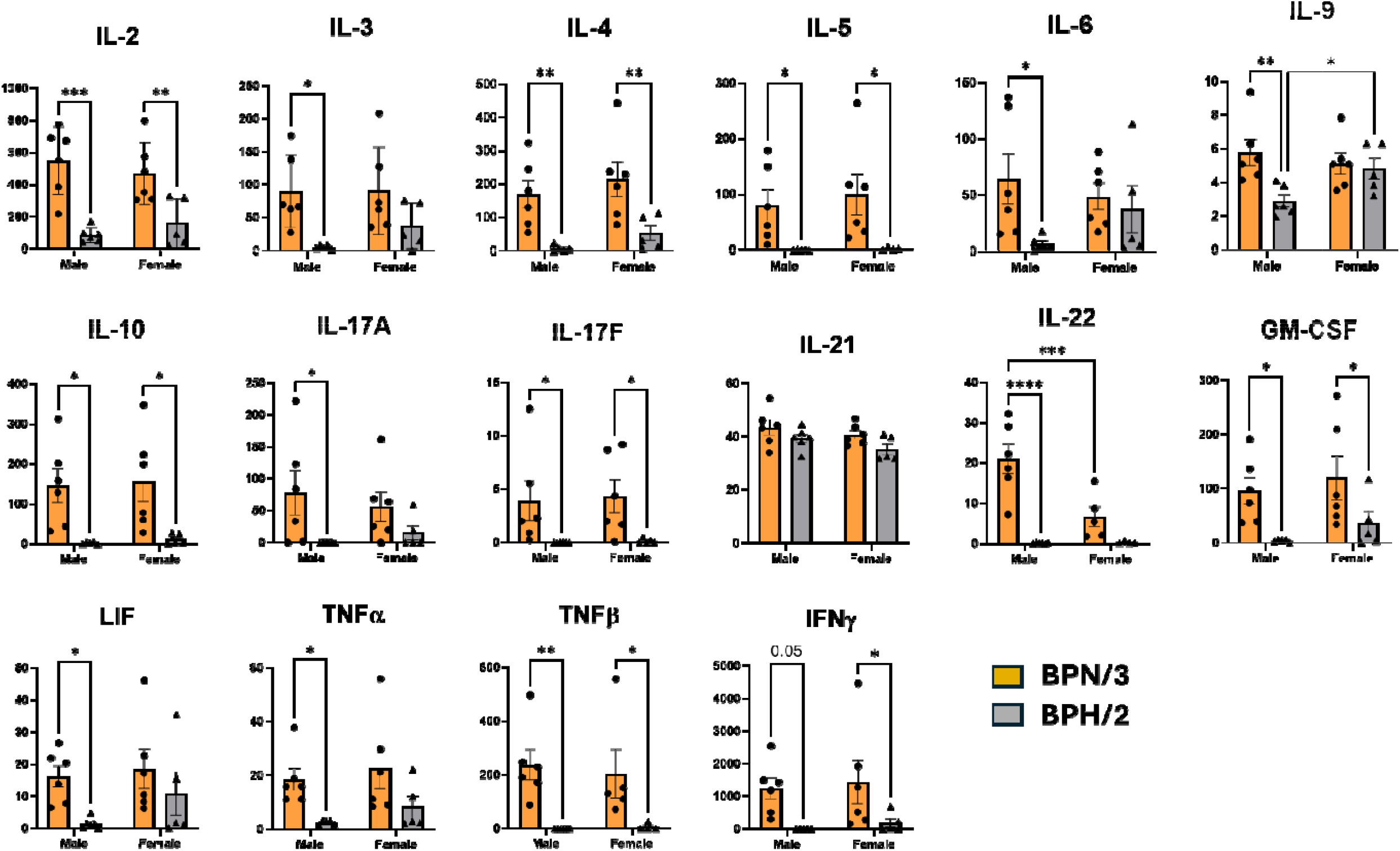
Decreased induction of T cell cytokines by splenic T cells from male and to a lesser extent, female, BPH/2 mice 120 h after activation. Freshly isolated splenocytes were cultured in the presence or absence anti-CD3/anti-CD28 activating antibodies. Cell supernatants were collected after 24 h for cytokine analysis by EVE Technologies (Calgary, Alberta, Canada). Data are presented as scatter plots plus bar graphs and include the mean ± SEM, n = 6. * p<0.05, ** p<0.01, ***p<0.001 and ****p<0.0001 as determined by 2-way ANOVA followed by Dunnett’s post hoc analysis.

## Discussion

The goal of this study was to understand whether there were fundamental differences between the immune systems of the hypertensive BPH/2 mouse model and the normotensive BPN/3 model. In addition, we wanted to determine whether there were sex differences between these two models. As expected, both male and female BPH/2 mice had significantly elevated mean arterial pressure as compared to BPN/3 mice, which was consistent with previous studies [19]. We were able to detect differences in blood pressure as early as 10 weeks and these were sustained up to 24 weeks of age. It is quite possible that the BPH/2 mice had increased blood pressure earlier than 10 weeks, but we are limited in how early we can measure blood pressure in part because the animals are acquired commercially. Thus, the age at which we can first start acquiring blood pressure data is determined by 1. the inability to order animals younger than 5-weeks of age in this strain, 2. the requirement for at least one week of acclimation prior to training mice for tail cuff measurements and 3. that it often takes several weeks of training before we can start to acquire tail cuff data. It should also be noted that in addition to blood pressure assessment by tail cuff measurement, we also attempted to measure blood pressures by radiotelemetry in a small subset of animals for this study. We were unsuccessful in this effort largely as a result of the small body mass of the BPH/2 female mice, which complicated the surgical implantation of the telemeters into these animals and who for the most part did not survive these surgeries.

An interesting observation from these studies was that the female BPH/2 mice had a greater mPVAT-to-body weight ratio, indicating a disproportionately larger mPVAT mass relative to body size in these animals. This was an unexpected finding because both the male and female BPH/2 mice are generally leaner than the BPH/3 mice overall. At this point, it is not clear why female BPH/2 mice have enlarged mPVAT. In future follow-up studies we would like to assess whether there is an increase in the number of adipocytes or adipocyte hypertrophy.

A major goal of these studies was to determine whether there were differences in immune cell function between male and female BPH/2 and BPN/3 mice. We assessed this by isolating immune cells from spleen and activating them ex vivo with a polyclonal T cell activator (anti-CD3/anti-CD28). The data clearly indicate marked differences between the two strains which are most pronounced in the male mice. Specifically, we found that cells derived from male BPH/2 mice showed a consistently dampened induction of nearly all T cell cytokines at both early (24 h) and late (120 h) time points after activation as compared to cells from male BPN/3 mice. There were only two exceptions to this trend in which neither IL-9 nor IL-21 differed between strains or sexes at 24 h and IL-21 was not different between groups at 120 h. Consistent with this, we also observed an attenuated induction of chemokines and secondary cytokines produced by myeloid and stromal cells in the male BPH/2 group. Overall, the data indicate a general hyporesponsiveness in the T cells derived from male BPH/2 mice. Although the underlying mechanism is not yet clear, there are multiple scenarios which could cause this effect. For example, it is possible that there is a defect in the machinery necessary for activation in T cells from male BPH/2 mice or an impairment in differentiation to an effector phenotype. It is also possible that the hyporesponsiveness is due to heightened T cell activity over time in male BPH/2 mice that resulted in T cell exhaustion. We are planning follow-up studies to differentiate between these possibilities.

Sex was a major factor determining the magnitude of immune cell responsiveness between the BPH/2 and BPN/3 strains in which the strain differences were much more pronounced in male mice. The sex differences were most noticeable at the early time point in which we observed diminished induction of 14 of the 16 T cell cytokines we measured in the male BPH/2 group in comparison to a decreased induction of only 5 cytokines in the T cells from the female BPH/2 mice. The sex differences waned over time, however. At 120 h after activation, there was a decreased induction of 15 of 16 cytokines in the male BPH/2 group as compared to decreased induction of 8 cytokines in the female BPH/2 group. Part of the sex differences can be attributed to differences in the magnitude of cytokine induction between the male and female BPN/3 groups. Specifically, we found that the induction of 7 out of 16 T cell cytokines was significantly decreased in the female BPN/3 group as compared to the male BPN/3 group. Furthermore, there was a nonsignificant trend toward decreased induction in another 4 – 5 cytokines in immune cells from female vs. male BPN/3 mice. The differences between the male and female BPN/3 groups seemed to equalize over time. At 120 h after activation, we observed diminished induction of only 2 cytokines in female BPN/3 group as compared to male. Taken together, the data suggest that immune cells from BPN/3 male mice show increased responsiveness to polyclonal activation in comparison to immune cells from both male BPH/2 mice as well as female BPN/3 mice. While the underlying mechanisms are not clear, the reasons could be related to differences in strength of T cell activation, efficiency of differentiation into an effector phenotype or some other factor (similar to possible reasons for differences in responsiveness between the male BPH/2 vs. BPN/3 strains stated above).

This study investigated differences in immunity between male and female hypertensive BPH/2 mice and normotensive BPN/3 control mice. The results indicate that in male mice, T cells from BPH/2 mice were markedly hyporesponsive to polyclonal T cell stimulation as compared to those from BPN/3 mice. These strain differences were much less pronounced in female mice. Overall, the results indicate markedly attenuated cytokine induction in immune cells from male BPH/2 mice. While the physiological consequence of these findings is not yet clear, the differences are striking and could ultimately impact immunity and inflammation.

